# Prosystemin–derived signals: bridging leaf microbiome dynamics and defense activation

**DOI:** 10.1101/2025.04.05.646382

**Authors:** Valeria Castaldi, Wisnu Adi Wicaksono, Francesca De Filippis, Gabriele Berg, Martina Chiara Criscuolo, Rosa Rao

## Abstract

The use of plant-derived peptides as resistance inducers provides innovative and potentially environmentally friendly methods to safeguard crop health. Although their use is increasing rapidly, their broader impact, particularly on plant-associated microbiomes, remains underexplored. This study investigated the influence of a promising immunomodulatory peptide derived from the tomato defense protein prosystemin on the tomato phyllosphere microbiome.

We applied the peptide via foliar spray biweekly to simulate common agricultural practices from the planting stage to two months post-germination. Using a shotgun metagenomics approach combined with qPCR, we identified bacterial communities of high abundance (up to 4.6 log_10_ bacterial 16S rRNA copies) and high diversity, mainly comprising Actino-, Alphaproteo– and Gammaproteobacteria, on all tomato leaves. The peptide treatment led to a significant and targeted shift in the bacterial community, characterized by reduced diversity and network complexity and species loss, i.e., *Streptomyces*. The enrichment was predominantly observed in bacterial genera such as *Acinetobacter*, *Sphingobium*, *Sphingomonas*, *Brevundimonas*, and *Massilia*, which are typically associated with improved plant growth and stress resilience. Intriguingly, shifts in both taxonomic and functional profile upon peptide application aligned with patterns typically observed during plant defense activation, involving jasmonic acid and related secondary metabolites. Members of the *Sphingomonadaceae* family, particularly *Sphingobium yanoikuyae*, have emerged as potential drivers of such microbial dynamics, likely adapting to plant upregulated defenses and potentially supporting its resilient phenotype.

Overall, in addition to its well-established role in combating tomato pests and necrotrophic fungi, the prosystemin-derived peptide paves the way for disentangling peptide-induced resistance and its interplay with the plant microbiota.

## Background

Agricultural practices aimed at protecting plants against abiotic and biotic stresses and promoting growth inevitably carry associated costs. Indeed, the use of traditional chemical pesticides often disturb non-target organisms and influence the balance of ecological processes. Addressing these challenges requires a comprehensive approach that integrates multiple strategies to safeguard agricultural productivity, ensure food security, and minimize health and environmental risks [1]. Building on the inherent ability of plants to cope with stress represents an innovative approach, aligning with processes already occurring in nature while offering sustainable solutions for plant health management [2–4]. For instance, there is a growing interest and investment in plant-derived peptide-based technologies due to their multiple capabilities that range from defense to biostimulation [5, 6]. Some peptides directly target pests, displaying antimicrobial, insecticidal, or nematicidal properties [6–8], while others, known as phytocytokines, primarily function in signalling and cell-to-cell communication, promoting immunity or growth [9–12]. Tomato Prosystemin (ProSys) has proven to be a valuable source for developing designed signaling peptides [13, 14], as it works as an hub protein containing bioactive motifs able to coordinate gene responses under different environmental challenges, thus counteracting a wide range of pests [15, 16]. However, the plant holobiont perspective has significantly expanded the way a resilient or resistant plant phenotype is defined [17, 18], underscoring how it is shaped by the interplay between plant genetics and its coevolved microbiome [19–21]. In this scenario, developing innovative plant protection strategies requires also a deep understanding of the dynamic interplay within the entire plant holobiont. The composition of the plant microbiota varies for each plant compartment as well as during a plant’s life cycle and is vertically transmitted and horizontally influenced [22]. The phytobiome community members indeed support plant health by producing bioactive molecules to control phytopathogens [23, 24], increasing nutrient uptake [25–27], inducing phytohormone production [28, 29], promoting germination and growth [30, 31], and degrading hazardous compounds, either in the air or soil compartments [32, 33]. This symbiotic functional interplay can be explained by plant‒microbe coevolution, and the plant genotype has been shown to be one of the most important drivers [34]. Considering these aspects, the application of signaling peptides to the host plant suggests the potential to influence the ecological dynamics of its interactions, potentially enriching or favoring microbial communities better adapted to specific physiological shifts [35–38]. Our prior work revealed that ProSys-derived peptides can mimic biotic stress, eliciting a primed defensive state in tomato plants by upregulating genes related to the JA pathway [13, 14]. This response confers a resistant phenotype aboveground, enabling plants to cope effectively with herbivorous insects such as *Spodoptera littoralis* larvae and necrotrophic fungi such as *Botrytis cinerea* and *Alternaria alternata* [13, 14, 39]. Here, we hypothesized that applying ProSys-derived peptides could also guide leaf-associated microbial communities, potentially shaping bacterial assemblages linked to improved plant resilience [20, 40, 41]. To shed light on this topic, we used a shotgun metagenomic approach to investigate the effects of a specific ProSys-derived peptide termed G [13] on the tomato phyllosphere microbiome, comparing it to those of control plants treated with PBS 0.1X solution (CTRL) or left untreated (UNT). We specifically explored the leaf phylloplane as a primary interface of exogenous agricultural treatment and a relatively underexplored compartment niche [42].

## Results

### Prosystemin peptide reduces bacterial diversity but selects specific taxa

We obtained a total of 768,712,768 high-quality reads that after host genome removal resulted in 19,605,952 high-quality bacterial reads (Additional file 1: Table S1) and 9,179 bacterial species across all samples. Compared with the CTRL and UNT treatments, the peptide treatment significantly altered the overall bacterial community structure at the species level (Fig. 1A) (*R^2^* = 0.575; *P* = 0.001). This pattern was related to a remarkable reduction in species richness (*P* = 0.002) and diversity (*P* < 0.001), which also appeared to be unevenly distributed (*P* < 0.001) according to the alpha diversity metrics (Fig. 1B). Our data suggest that under the conditions imposed by peptide application, selective pressure occurs, favoring certain bacterial taxa, which leads to an increase in abundance and dominance over other taxa.

**Figure 1.**
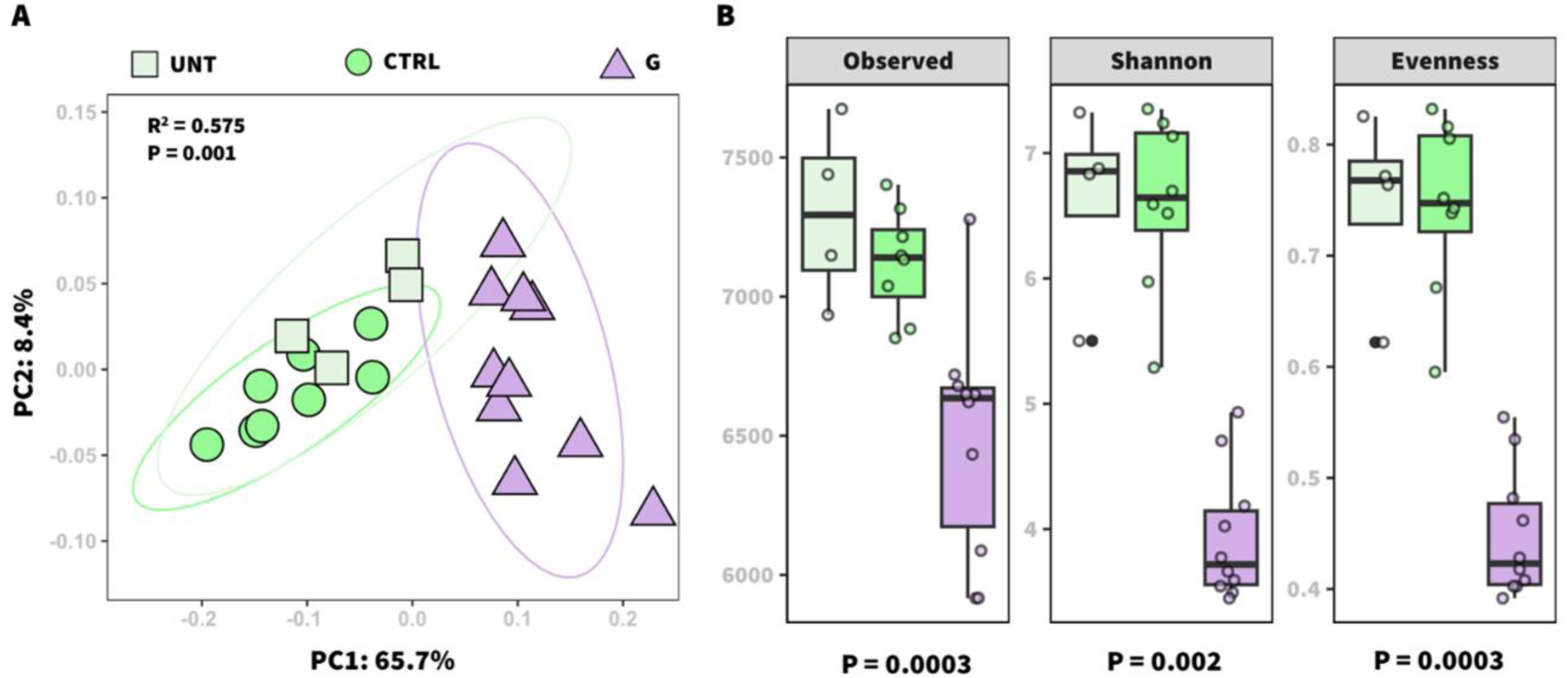
Bacterial structure and diversity of the tomato leaf epiphytic community in untreated, control (PBS 0.1X), and G peptide-treated (100 fM) plants. Bray–Curtis distance matrices of bacterial community structures between samples were visualized using a two-dimensional PCoA plot (A). The biological significance of the samples was tested via PERMANOVA. Richness (observed species), Shannon, and Pielou’s evenness indices (B) were calculated to explore the overall alpha diversity within the samples. The Kruskal‒Wallis test was used to test the significance of the data. UNT: untreated, CTRL: control-, G: peptide-treated plants.

### Alphaproteobacteria and Gammaproteobacteria dominate the peptide-treated leaves

The taxonomic composition at the class level of the leaf microbiome (Fig. 2A) revealed that, compared with the CTRL (32.7% and 15.9%) and UNT (29.7% and 16.9%) plants, the G peptide favored the multiplication of Alphaproteobacteria and Gammaproteobacteria (48.3% and 31.8%), respectively. In contrast, Actinomycetes predominated in both the control plants (33%) and the peptide-treated plants (10.1%). Given the negligible differences in the observed, Shannon, and beta diversity metrics (*P* = 0.08; *P* = 1; *P* = 0.1) between the PBS-treated and untreated plants, the analyses were conducted with a focus on comparing the G-treated and PBS-treated plants. The differential abundance analysis conducted with edgeR identified a total of 1,612 bacterial species with a significant positive log_2_-fold-change (log_2_FC) and 933 with negative log_2_FC in G-treated samples (Additional file 1: Table S2). Bacterial species considered enriched or depleted (log_2_FC > 2 or < −1, *P* < 0.05, with relative abundance > 0.05%) were retained and are shown in Figure 2B.

**Figure 2.**
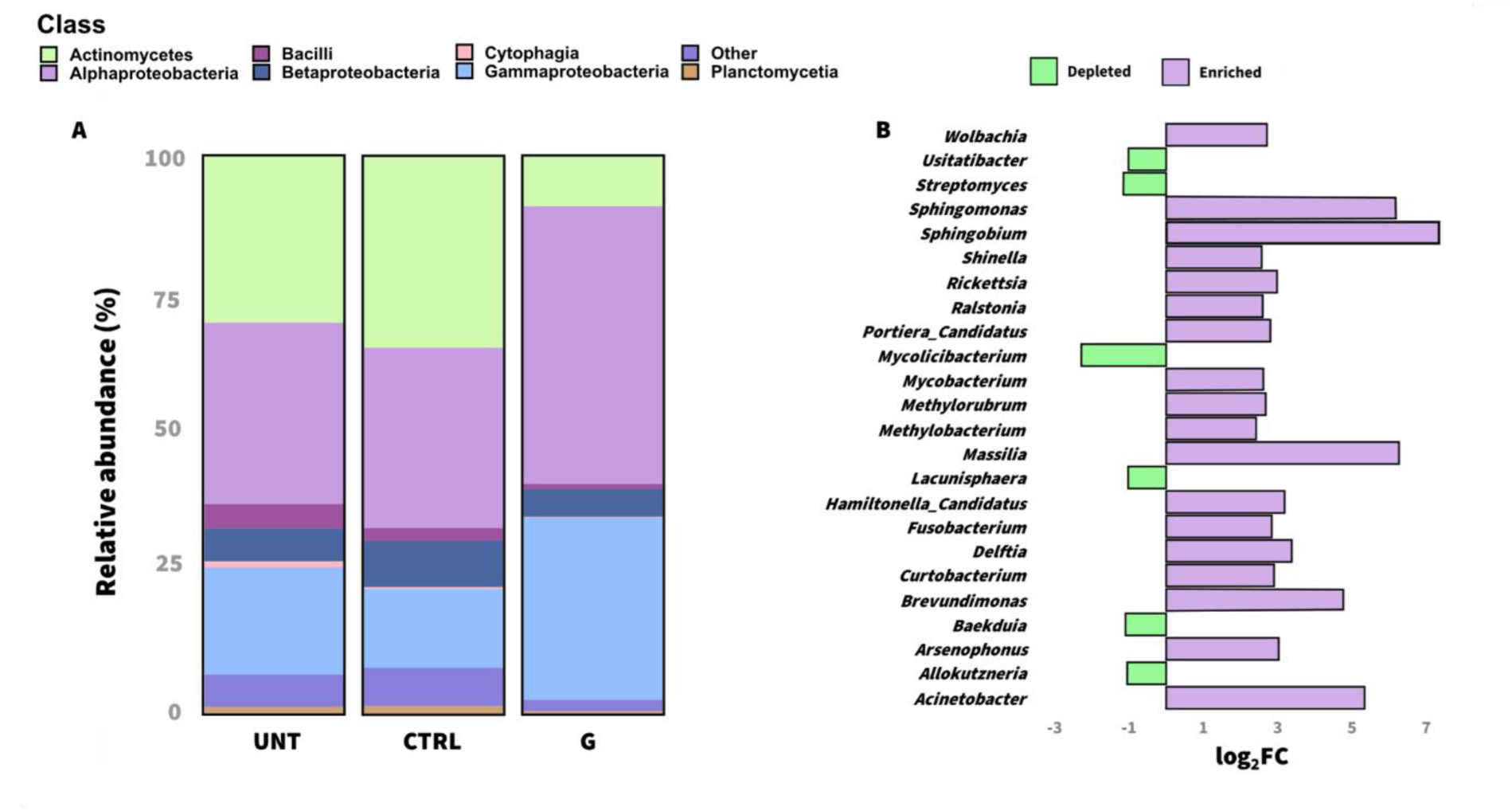
Bacterial community composition of *S. lycopersicum* cv. San Marzano Nano leaves subjected to different foliar spray treatments. Mean relative abundance of bacteria at the class level (**A**) in tomato plants under three different treatments: UNT (untreated), CTRL (0.1X PBS), and G peptide (100 fM). Differential abundance analysis (**B**) performed at the bacterial genus level in the G-treated plants compared with the CTRL plants via the edgeR package. Positive log_2_-fold-change (log_2_FC) values (violet bars) indicate higher abundances of the respective bacterial genera in the G-treated samples, whereas negative values (green bars) represent higher abundances in the PBS-treated plants. Significantly enriched or depleted bacteria (*P_adjusted_* < 0.05) with a relative abundance > 0.05% are shown.

Among the Alphaproteobacteria, enrichment of genera such as *Brevundimonas* (log_2_FC = 4.0, *P* < 0.001), *Sphingobium* (log_2_FC = 6.28, *P* < 0.001) and *Sphingomonas* (log_2_FC = 4.63, *P* < 0.001) were observed, whereas among *Gammaproteobacteria* and Betaproteobacteria, *Massilia* (log_2_FC = 6.27, *P* < 0.001) and *Acinetobacter* (log_2_FC = 5.03, *P* < 0.001) were the affected bacterial genera. Specifically, at the bacterial species level, *Sphingobium yanoikuyae* (log_2_FC = 7.33), *Sphingomonas hankookensis* (log_2_FC = 6.13) and *Acinetobacter johnsonii* (log_2_FC = 4.35) demonstrated significant enrichment due to the peptide treatment (Fig. 3A) and were among the core bacterial species associated with this treatment (Fig. 3B). Surprisingly, we also found that the abundances of some bacterial genera representative of insect endosymbionts increased after treatment (Fig. 2A). Among them, *Rickettsia* (log_2_FC = 2.93, *P* < 0.001) and Hamiltonella_Candidatus (log_2_FC = 3.18, *P* < 0.001) may influence plant behavior during stressful events. In the PBS-treated plants, *Streptomyces*, classified within the *Actinomycetes* class, was among the most abundant genera (9,23%), remarkably affected by peptide application (1.94%). Moreover, most *Streptomyces* spp. constitute the core of both control conditions (Fig. 3B; Additional file 1: Table S3).

**Figure 3.**
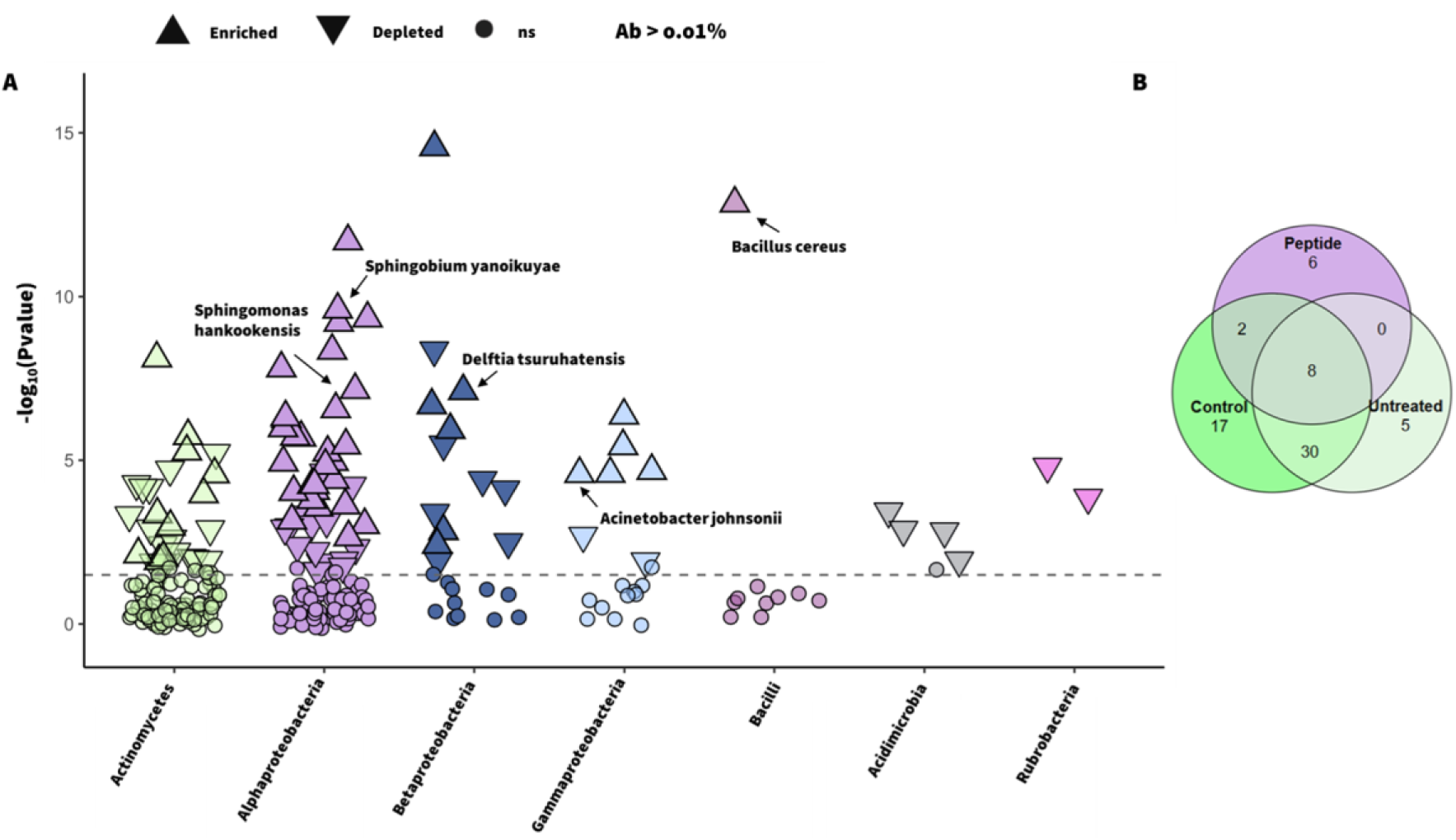
Enriched bacterial species in the G treatment group compared with those in the CTRL group and a Venn diagram showing the core and unique number of species among the treatment groups. Manhattan plot (**A**) showing enriched (upward-pointing filled triangles), depleted (downward-pointing filled empty triangles), and nonsignificant (ns, filled circles) bacterial species in the peptide-treated plants compared with the control (PBS). Species were filtered by *P* < 0.05 and a relative abundance (Ab) > 0.01%. Shared and unique core bacterial species among the three treatments (**B**) are shown with a Venn diagram. Core species were filtered by prevalence (75%) across samples and Ab > 0.1%

### qPCR reveals a higher bacterial load in G samples and strengthens the importance of few taxa

The absolute bacterial abundance in UNT and CTRL leaves, measured through quantitative polymerase chain reaction (qPCR), was 4.12 ± 0.4 and 4.37 ± 0.5 log_10_ bacterial 16S rRNA gene copies, respectively (Fig. 4). Whereas leaves treated with G peptide showed a higher bacterial load (4.60 ± 0.4 log_10_ bacterial 16S rRNA gene copies) compared to untreated leaves. Although not statistically significant, a similar trend was observed for Alphaproteobacteria and *Sphingomonas*, with a higher abundance in G peptide-treated leaves (4.22 ± 0.3 log_10_ and 3.28 ± 0.3 log_10_ copies, respectively) relative to UNT (3.82 ± 0.5 log_10_ and 2.88 ± 0.4 log_10_ copies) and CTRL leaves (4.03 ± 0.2 log_10_ and 3.10 ± 0.2 log_10_ copies).

**Figure 4.**
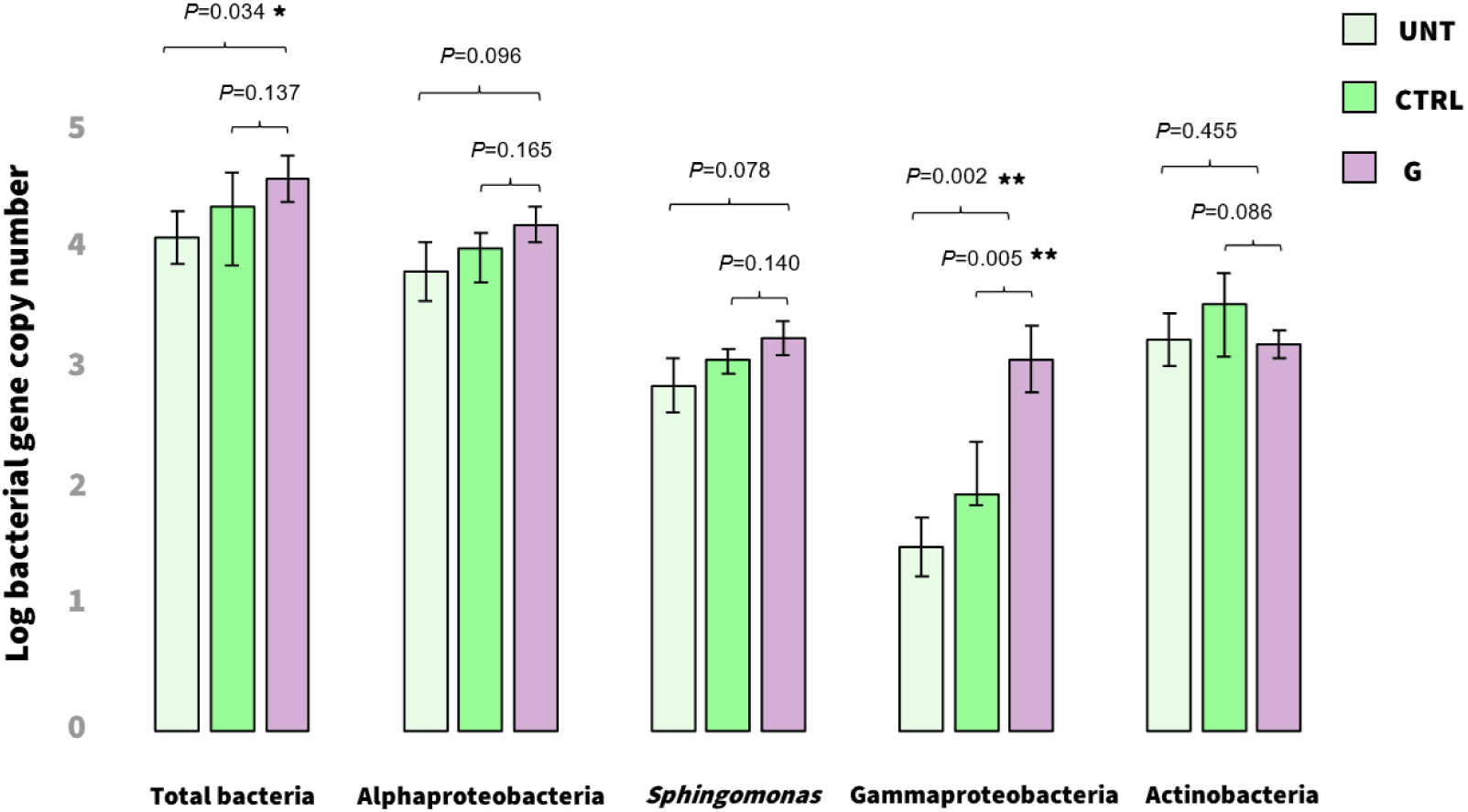
Absolute abundances of total bacteria, Alphaproteobacteria, *Sphingomonas*, Gammaproteobacteria and Actinobacteria calculated with quantitative real-time PCR (qPCR) between untreated (UNT), PBS 0.1X-treated (CTRL), or peptide-treated (G) tomato leaves. Statistical significance was assessed by Kruskal–Wallis with Dunn’s test for multiple comparisons (\**P* < 0.05, \*\**P* < 0.01). Error bars indicate the standard error.

Notably, G peptide treatment led to a significant enrichment of Gammaproteobacteria (3.10 ± 0.5 log_10_ Gammaproteobacteria copies) compared to untreated (UNT; 1.53 ± 0.5 log_10_) and control conditions (1.97 ± 0.8 log_10_). In contrast, Actinobacteria were more abundant in control leaves than in those treated with G peptide or left untreated. These data further support the trends already observed with shotgun metagenomic data analyses following G peptide treatment.

### The peptide redefines the interaction balance on the phylloplane toward a simple network

To explore the interactions among the bacterial community, two intra-kingdom co-occurrence networks were constructed to compare the CTRL– and G-associated leaf communities. In both networks (*r* = 0.7, *P* < 0.05), the nodes represent the 400 most abundant bacterial species, but a notable difference in edge numbers between the two conditions was observed. The control (on the left) had a greater proportion of edges (41,438) and a shorter average distance between nodes (1.73), indicating a denser structure with both positive (94.65%) and negative (5.35%) correlations across a wide range of bacterial species (Fig. 5A-B). In contrast, peptide treatment (on the right) led to a sparser network with fewer (15,995) and more negative (6.63%) but selected connections. Interestingly, G results in greater modularity (0.47 vs. 0.09), indicating a stronger community structure with more distinct modules (7 vs. 4) (Additional file 1: Table S4). This suggests possible ecological patterns where bacterial species belonging to the *Actinomycetes* and *Proteobacteria* classes tend to group together within their respective classes but remain distinct from each other (Fig. 5B). The level of complexity was measured on the basis of the average degree (> 60) and closeness centrality (> 0.4), revealing the presence of more bacterial hubs, highly connected and influential, in the CTRL than in the peptide-treated plants (Fig. 5C). However, despite their reduction, certain genera (e.g., *Acinetobacter*, *Brevundimonas*, and *Massilia*; Fig. 5D) presented more connections than the CTRL plants, potentially playing key roles in the reshaped community.

**Figure 5.**
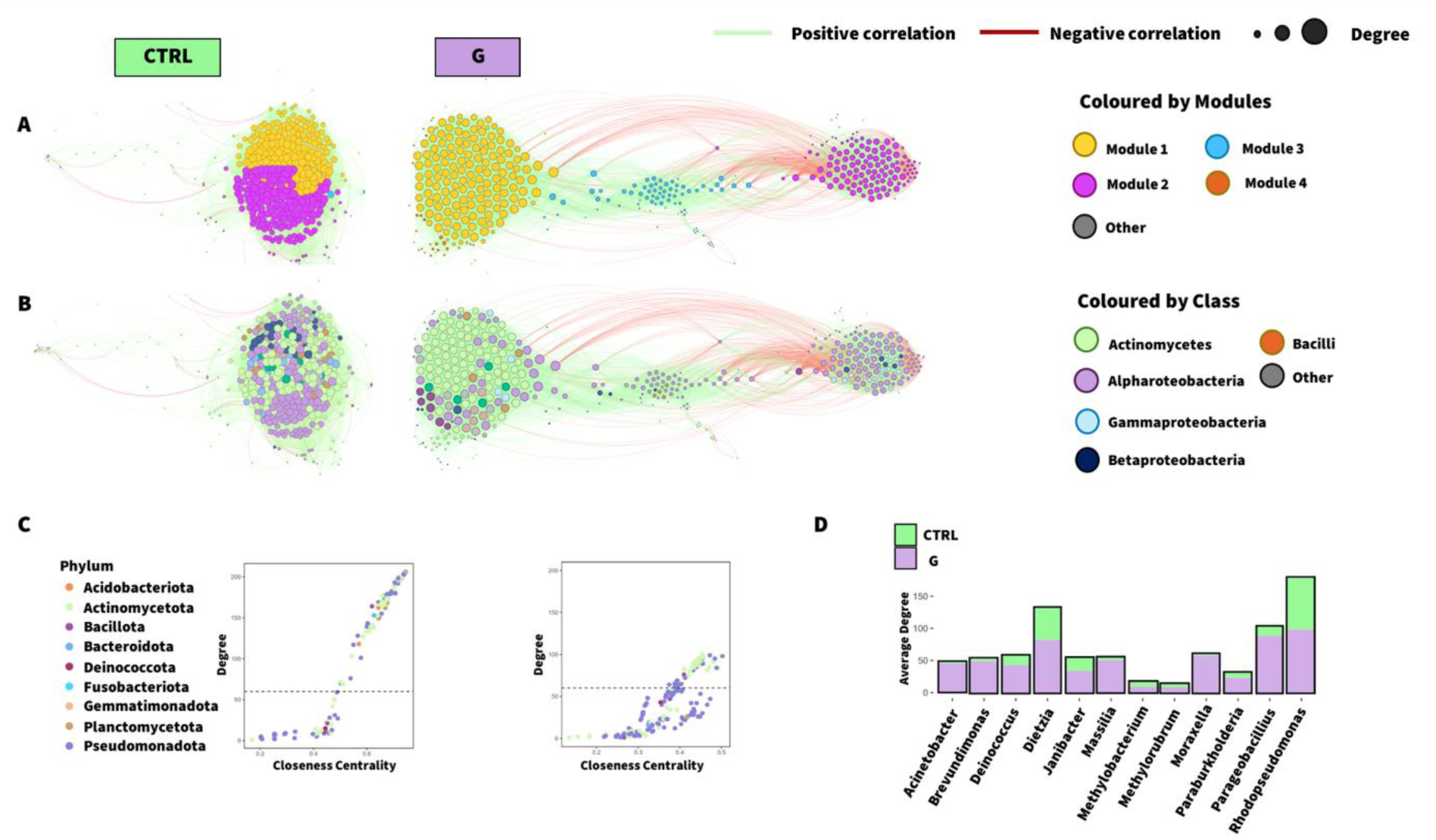
Bacterial intra-kingdom co-occurrence networks in the control (left side) and ProSys-derived G peptide (right side)-treated tomato leaves. Each node represents the 400 most abundant bacterial species, which are colored according to module (**A**) and class taxonomic rank (**B**). Positive and negative correlations are shown by green and red edges, respectively. Scatter plots (**C**), colored by phylum classification, show the hub node distribution according to degree and closeness centrality scores. The bacterial genera that gained more connections on the basis of average degree scores (**D**) are shown in violet for the G treatment and in green for the CTRL treatment. The Kruskal‒Wallis test was used to test the significance of the difference in network connectivity at the genus level (χ² = 45.094, *P* < 0.001).

### Functional profiling reveals intense bacterial activity to cope with changes in the environment

Our analysis of the bacterial community structure led us to explore its potential metabolic properties and ecological functions in greater depth. High-quality filtered contigs were clustered together to identify potential protein-coding regions (ORFs). Among the 381,825 predicted ORFs, 189,644 were successfully annotated, with 56,7% of these genes assigned to KEGG Orthology terms (KEGG-KO). Beta diversity analysis (Fig. 6A) confirmed that the observed shift in the bacterial community composition was further explained by functional differences (*R^2^* = 0.366, *P* = 0.001). According to KEGG-KO enrichment analysis performed via both the *ClusterProfiler* and *EdgeR packages* (log_2_FC > 2, *P_ajdusted_* < 0.05) on the G-treated samples, the functional shift was attributed to increased metabolic processes and responses to the environment (Fig. 6B).

**Figure 6.**
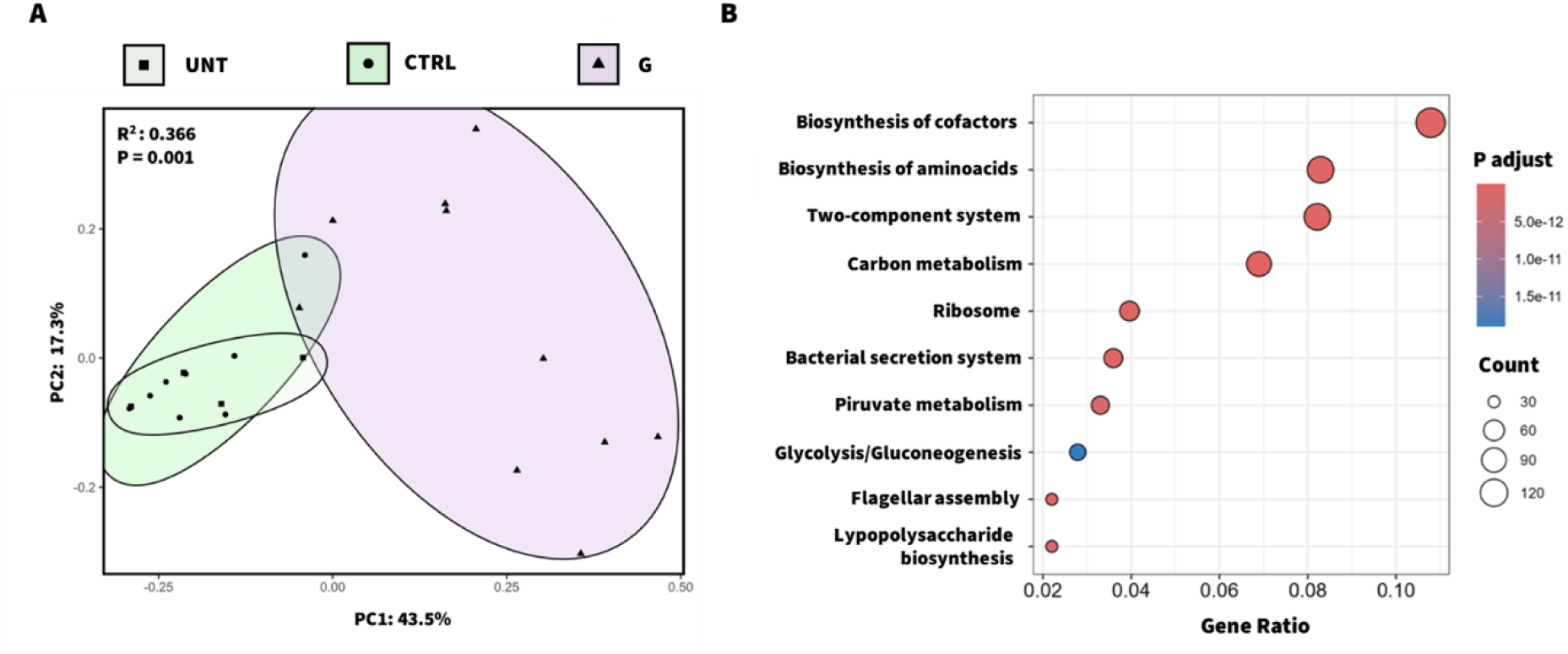
Functional exploration of the leaf-associated bacterial community. A Bray‒ Curtis matrix (**A**) was used to visually explore the differences between the genes of the phyllosphere samples according to the different treatments. Top 10 KEGG pathways (**B**) based on the KEGG Orthology terms (KOs) associated with the enriched genes of the peptide-treated samples. The circle sizes represent the number of enriched-KOs grouped according to KEGG pathways (third level). The analysis was performed by the Microbiome Profiler package in R.

Specifically, we observed an increase in the abundance of genes associated with the biosynthesis of cofactors (147 KOs) and the synthesis of vitamins, including B12 (M00122, M00924, M00925), B6 (M00124) and B7 (M00123), in plants treated with the G peptide, which may support both bacterial and plant performance. Additionally, the biosynthesis of essential cofactors associated with energy production and redox balance, such as tetrahydrofolate (M00841, M00842), coenzyme A (M00120), coenzyme Q (M00117), and molybdenum (M00880), was also enriched. The increase in the abundance of genes associated with amino acid biosynthesis (113 KOs) suggests that the bacterial community may participate in nutrient cycles or provide precursors for the synthesis of plant compounds, therefore influencing plant productivity and resilience. For example, we found that entirely represented modules for the synthesis of key amino acids, such as tryptophan (M00023), lysine (M00016), proline (M00015) and methionine (M00017), were enriched in plants treated with the G peptide. These amino acids may be potentially supplied to the plant by the associated microbiome. Notably, the observed enrichment of two-component system (TCS) (112 KOs) and bacterial secretion system (49 KOs), indicates that the community is highly responsive to environmental signals. These signals may originate not only from other bacteria competing to survive under newly established phyllosphere conditions but also from the plant itself following G peptide application.

### TCS reveals potential mechanisms of interaction with the plant host

The two-component system is a crucial pathway that bacteria use to interact with the environment. Therefore, understanding how bacterial performance and plant responses are influenced by external factors, such as peptide application, is key. By filtering the eggNOG gene table for TCS-associated KEGG terms, we performed a beta diversity analysis (Additional file 2: Fig. S1A), which revealed that the G-treated samples formed a distinct cluster from the PBS-treated plants (*R^2^* = 0.300; *P* = 0.001). Indeed, the G-treated samples presented significant enrichment (n = 175) rather than depletion (n = 10) of differentially abundant genes (log_2_FC > 2 & < –1; *FDR* < 0.05; EdgeR) associated with relevant KEGG categories in comparison with the PBS-treated plants. We identified an increased abundance of genes associated with secretion and extracellular structure formation (Additional file 2: Fig. S1B; Additional file 1: Table S6). For instance, an enrichment of *wza* (K01991), a polysaccharide biosynthesis export protein, and *TolC* (K12340), a component of the type I secretion system involved in the transport of enzymes and toxins to the extracellular environment. Additionally, the endoglucanase gene *egl* (K01179), which is involved in the degradation of polysaccharides such as cellulose, was more abundant (log_2_FC = 5.45), suggesting a role in bacterial colonization of leaves or nutrient recycling. The data also highlighted enriched gene sets associated with chemotaxis (*mcp*, *cheA*, *cheW*, *cheR*, *cheB*: K03406, K03407, K03408, K00575, K13924), quorum sensing (*qseB*, *qseC*: K07666, K07645) and biofilm formation (*envZ*, *ompR*, *ompF*: K07638, K07659, K09476). These findings suggest that bacterial reorganization and adaptation to new leaf surface conditions are critical for successful colonization and competition within the phyllosphere environment. Moreover, an enrichment of genes potentially involved in plant nutrition, particularly those associated with phosphate assimilation and uptake, such as *phoA*, *phoB*, and *phoR* (K01077, K07657, K07636), was observed. The overall enriched genes were predominantly associated with *Alphaproteobacteria* and *Gammaproteobacteria*, with *Sphingomonadaceae* (42%) and *Moraxellaceae* (28%) being the most represented families.

### Sphingobium yanoikuyae: beyond bioremediation

Among 47 metagenome-assembled genomes (MAGs) with completeness over 50% and contamination below 10% (Additional file 1: Table S7), *Sphingobium yanoikuyae* (MAG ID: G_10) and *Acinetobacter johnsonii* (G_1) were the most significantly enriched MAGs in the G peptide-treated samples (Additional file 2: Figure S2; Additional file 1: Table S8).

To further explore the potential plant growth-promoting traits (PGPTs) within the microbial communities promoted by the peptide, we used the PGPT-Pred tool, part of the PLaBAse web resource [43]. We focused on high-quality, dereplicated MAGs (completeness >90% and contamination <5%) corresponding to bacterial species enriched by G peptide treatment for a more detailed overview of potential PGPT-related ORFs. The analysis revealed that *Sphingobium yanoikuyae* (MAG ID: G_10, n=4) harbored the greatest number of PGPT-associated genes (1,398), followed by *Brevundimonas* (G_9, n=4), with 1,136, and *Rickettsia* (MAG G_3, n=16), with 443 predicted genes. *Sphingobium yanoikuyae* is commonly associated with bioremediation abilities, such as detoxification of heavy metals or xenobiotics. Accordingly, PGPT-Pred tool assigned 115 genes to “heavy metal detoxification” and 73 to “xenobiotics biodegradation”, compared to *Brevundimonas* (91 and 35) or *Rickettsia* (30 and 13), respectively (Additional file 1: Table S9).

However, *S. yanoikuyae* and *Brevundimonas* presented similar profiles, with genes involved in plant colonization (26%), stress-related biocontrol (20% and 21%, respectively), and plant nutrition (14%) (Fig. 7; Additional file 1: Table S9).

**Figure 7.**
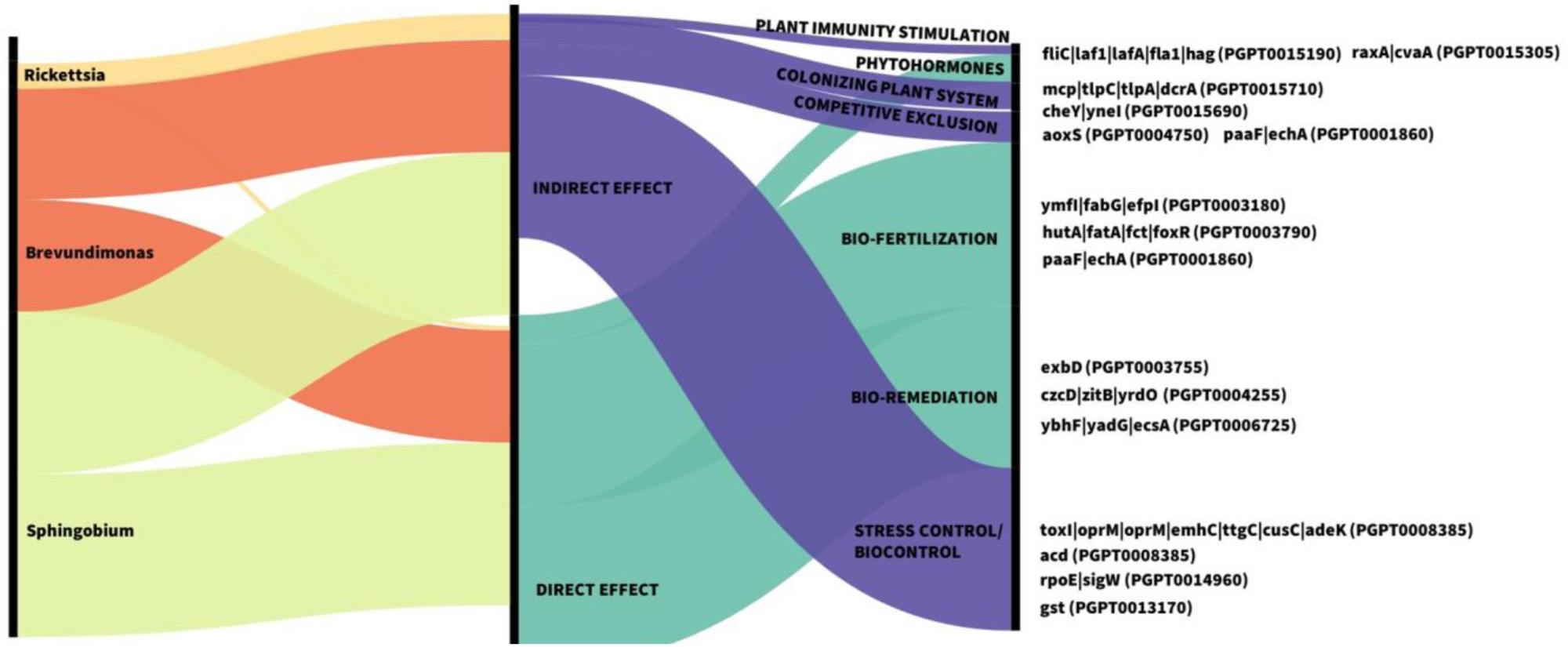
Peptide-associated MAGs and their plant beneficial traits (PGPTs) according to PLaBAse. Alluvional diagram showing MAGs associated with peptide-treated plants and classified into the *Rickettsia, Brevundimonas, and Sphingobium* genera. The bacteria represent the source connected to different levels of plant beneficial trait classification according to PLaBAse PGPT-Pred tool. The figure shows level 1 (direct and indirect effects), level 2 (plant immunity stimulation, colonizing the plant system, competitive exclusion, bio-fertilization, bioremediation, and biocontrol), and level 6, with the details of some of the genes assigned to the PGPT categories. Different colors are used to represent each level, and the thickness of the connections between levels is proportional to the frequency of genes within each category, indicating the abundance of each trait.

Notably, *S. yanoikuyae* showed the highest gene count (n=75) for *hutA, fatA, fct*, and *foxR* (PGPT0003790), associated with the TonB-dependent receptor (K02014), which plays a key role in iron uptake, a crucial strategy for counteracting biotic stressors such as fungal pathogens. These findings suggest that *S. yanoikuyae* may have multiple functional roles, particularly in stress mitigation, beyond its previously recognized functions in bioremediation [44–46].

## Discussion

Compared to control conditions, treatment with the bioactive 16-amino acid signaling peptide [13] induced a significant shift in the epiphytic phyllosphere bacterial community of tomato plants, as indicated by increased beta diversity distance and a reduced alpha diversity. Although greater microbial diversity is often associated with a “protective” microbiome, its definition and consistency across plant species and environmental contexts remain challenging [47, 48]. For instance, gamma-aminobutyric acid (GABA), which confers resistance to pests and promotes VOC production, was shown to reduce microbial diversity while favoring a suppressive microbiome against *southern leaf blight* (SLB) [49]. Likewise, repeated foliar application with small peptides as fertilizers decreased bacterial richness in the tea phyllosphere but promoted the enrichment of microbial taxa known to promote plant growth and immunity [35]. These findings suggest that microbial diversity alone may not be a reliable indicator of a healthier microbiome; instead, the establishment of targeted and functionally specialized taxa should be considered a more relevant factor [47]. In our study, ProSys-derived G peptide promoted notable enrichment of Alphaproteobacteria, such as *Sphingomonas*, *Sphingobium*, and *Brevundimonas*, as well as Gammaproteobacteria, such as *Acinetobacter*, whereas some members of the Actinomycetes, including *Streptomyces*, were reduced.

Proteobacteria and Actinomycetes prominently shaped the community structure in G-treated leaves, characterized by high modularity and a greater proportion of negative interactions. In contrast, the CTRL condition resulted in a more cohesive, interconnected, and diverse community. These features typically reflect higher stability in both mutualistic and antagonistic interactions [50], whereas disruptions, such as those caused by intensified farming practices [51], tend to reduce network complexity.

Activation of plant defense pathways is also known to disrupt microbial networks [52]. Indeed, shifts in metabolic processes, including changes in root exudate composition, can lead to decreased microbial density and connectivity [53, 54]. Exogenous treatments may introduce minor disturbances within plant microhabitats [47, 55], promoting the survival of taxa that are better adapted to the selective pressures imposed by repeated peptide-induced plant responses. For example, application of RALF23, a signaling peptide involved in plant immunity, promoted the enrichment of fluorescent pseudomonads in the *Arabidopsis* rhizoplane, which successfully suppressed the pathogen *Fusarium oxysporum* [56]. Thus, the increased modularity and negative correlations observed in the G-treated samples may reflect a more compartmentalized network, potentially stabilizing interactions within selected microbial groups under new competitive conditions [52].

Our findings resemble observations in other plant species where JA-related defense activation similarly modulated microbial community dynamics. For example, JA-mediated defenses in wheat or rice are associated with reduced bacterial diversity in root microbiomes, whereas JA-deficient *Arabidopsis* mutants present increased diversity [57–59]. In addition to plant genetic modification, exogenous treatments with JA also reduce the Shannon evenness of the bittercress phyllosphere microbiome [60], supporting the idea that elicitor molecules, whether acting endogenously or exogenously, can similarly reshape the plant-associated microbiome.

Therefore, we speculate that the observed effects on taxa composition, interactions, and functionality are driven by plant defense metabolism triggered by the immunomodulatory effect of the peptide, which mimics external biotic stress [61]. Through the JA pathway, ProSys and its derivatives trigger the synthesis of secondary metabolites such as phenylpropanoids [14, 39, 62, 63], which can in turn affect the plant-associated microbiome [64]. Among glycoalkaloids, α-tomatine, known for its antifungal and pesticidal properties, is associated with JA overexpression [65, 66], MeJA treatment, and wounding. However, its levels are reduced in ProSys-impaired tomato mutants [61, 65]. In terms of the plant-associated microbiome, tomatine production has been linked to decreased bacterial Shannon diversity, lower abundance of specific Actinobacteria, and selective recruitment of taxa such as members of the *Sphingomonadaceae* family and the genus *Sphingobium* (e.g., *S. yanoikuyae*) [67–69]. These bacteria are capable of detoxifying α-tomatine and, in turn, contribute to host resistance [67–69]. Similarly, enrichment of *Sphingomonadaceae* was observed in the phyllosphere of bittercress under herbivory stress [60], while low-tomatine-producing tomato lines with impaired JA pathway showed greater susceptibility to *Spodoptera* attacks and reduced *Sphingomonadaceae* abundance [70]. As suggested by Tronson et al. [71], the specific contribution of this bacterial family under biotic stress conditions, such as herbivory, should be further investigated. We propose that the success of taxa enriched by peptide treatment may be related to their ability to cope with the plant’s primed defense state, potentially through biofilm formation or detoxification mechanisms, as suggested by our functional analyses. *Sphingomonas* and *Sphingobium*, for example, are well known for their ability to detoxify several compounds, including phenolics and glycoalkaloids, which act as pest deterrents in plants [68, 72, 73]. However, the significant reduction in Actinomycetes, particularly *Streptomyces*, in peptide-treated plants cannot be ignored, as these bacteria are widely recognized as valuable producers of secondary metabolites that promote plant growth and resilience against pathogens [59, 74]. They are found primarily in the roots and gain a competitive advantage, particularly under drought (associated with ABA production) or iron deficiency [75]. Previous studies reported a decrease in *Streptomyces* abundance in wheat following MeJA treatment [59] and an increase in the rhizosphere of *Arabidopsis* mutants defective in the JA pathway [76]. Intriguingly, scopoletin, a phytoalexin induced by MeJA [77], inhibited the growth and activity of the pathogen *Streptomyces scabiei* [64], a species closely related to *S. caniscabiei*, whose abundance was significantly reduced by peptide treatment in our study [78] (Additional file 1: Table S2). In contrast, low JA concentrations *in vitro* promoted the growth and antibiotic production of some *Streptomyces* species [79]. Given this perspective, we cannot definitively claim that the peptide treatment *enhanced* the microbiome, as the enrichment of certain beneficial species was accompanied by a reduction in others that also contribute positively to plant health. The treatment also affected bacterial biodiversity and connectivity, leading to shifts in the abundance of genera typically associated with the phyllosphere of vegetable crops, some of which may carry antibiotic resistance genes (ARGs) [80–82]. This underscores the need for further investigation of both genetic and environmental factors that influence plant preferences in hosting bacterial taxa potentially carrying ARGs, and how these shape microbial ecological niches [50, 82, 83]. The functional profiles associated with G application revealed increased bacterial motility and biofilm formation, similar to the effects of MeJA treatment in tomato plants [84, 85]. Among the most abundant genes, *wza* (K01991) is involved in the extrusion of exopolysaccharides (EPSs), which are crucial for biofilm formation [86, 87]. As part of the TCS signaling pathway (K02020), EPS and quorum sensing work as driving forces for biofilm formation, which not only is crucial for plant tissue colonization but also provides protection against salinity, drought, and pathogen attacks [88–90]. TCS, which is involved in the bacterial response to environmental stress [54, 91], along with the bacterial secretion system, has been previously linked to the beneficial roles of *Sphingomonas* and *Bacillus* since they are actively recruited via root exudates during *F. oxysporum* infection in cucumber plants [91]. Our KEGG-KO enrichment analysis further indicated that the peptide enhanced the biosynthesis of cofactors, vitamins, and amino acids. Intriguingly, similar functional enrichment was attributed to the rhizosphere microbiome of cowpea under herbivory stress caused by the leafminer *Liriomyza trifolii* [92]. Vitamins of the B group as well as amino acid production by plant-associated microbiota are linked to growth promotion, plant defense, and selective adaptation [24, 93–96]. Tryptophan (Trp), for example, serves as a precursor of the auxin indole-3-acetic acid (IAA), which promotes development and abiotic stress resistance [97]. Enhanced Trp biosynthesis is associated with JA-mediated responses against *Spodoptera littoralis* attacks [98], whereas its levels are diminished in the root exudates of plants with impaired JA production [76].

Among the genes enriched with the G peptide, we also identified *nif* and *fix* genes, which are essential for N fixation, and *nir* genes, which are involved in N assimilation and are associated mainly with the *Acinetobacter* and *Sphingobium* genera. Accordingly, during herbivory stress, plants may adopt a “cry-for-help” strategy for additional nitrogen reallocation for the synthesis of defense-related compounds. This is supported by the activity of *Rhizobiales* and *Sphingomonadales*, able to counteract the aboveground biotic threats in cowpea [92].

Notably, the enrichment of the iron complex outer membrane receptor protein *TonB* (K02014) in *Sphingobium yanoikuyae* in our study, was recently associated with the ability of *Sphingomonas* to suppress the necrotrophic pathogen *Diaporthe citri* in the citrus phyllosphere, through competitive iron acquisition [99]. *S. yanoikuyae*, which was isolated from the phyllosphere of the tropical plant *Dracaena marginata*, has also shown inhibitory activity against the fungal pathogen *B. cinerea* [100]. In addition to its well-established role in environmental bioremediation, *S. yanoikuyae* has been increasingly associated with growth improvement and biocontrol properties in recent years [101, 102], as further supported by the PLaBAse analysis in this study.

Overall, we speculate that members of *Sphingomonadaceae*, including *S. yanoiukaye*, may be indirectly selected by plant-activated defenses, ranging from phytohormones such as JA to key metabolites, due to their stress-coping capacities. In turn, their presence may confer adaptive benefits to plants under primed conditions.

## Conclusion

Our findings demonstrate that the application of ProSys-derived peptide resulted in a significant and targeted shift in the tomato leaf microbial community, affecting both the taxonomic composition and functional traits. This reshaping appears to be driven primarily by the plant defense response triggered by the peptide, which is in line with our previous research on the indirect biological effects of ProSys peptides on fungal pathogens and insect pests. The observed changes resemble those typically observed in plants with upregulated JA defenses or treated with its derivatives, particularly with the synthesis of specific secondary metabolites that may act as chemoattractants within quorum-sensing mechanisms activated by bacteria. Given this perspective, it will be crucial to understand the direct role of the enriched bacteria on tomato plants and their responses to their defense-related metabolites, elucidating (1) potential recruitment mechanisms on the leaves, (2) the ecological significance of microbial networks upon defense activation in a time-course analysis and (3) the microbiome contribution to pest control after priming with the peptide. Such studies will add valuable insights into the “cry-for-help” strategy among different plant compartments.

## Methods

### Plant material and growth conditions

In April 2023, *Solanum lycopersicum L.* cultivar “*San Marzano nano*” tomato seeds were germinated in a growth chamber at a controlled temperature of 24±1°C, 60±5% relative humidity (RH) and complete darkness. The seeds were placed in Petri dishes on moist, sterile paper for a few days until the emergence of the rootlets. The plantlets were subsequently moved to a polystyrene tray containing autoclaved soil mixed in a greenhouse growth chamber at 26±1°C with 60±5% RH and an 18:6 h light/dark photoperiod. Following an adaptation period of two weeks, the plants were transplanted into 9-cm-diameter pots filled with autoclaved soil mixture and kept under the same conditions until they reached two months of growth.

### Treatment experimental design

Two-week-old plants were subjected to four foliar spray treatments every 15 days, with volume amounts adjusted according to their growth stage and size. The experimental plan included the three following treatments: 100 fM ProSys-derived peptide (referred to as G) diluted in 0.1X phosphate-buffered saline (PBS) as previously reported [13], 0.1X simple PBS representing the control (CTRL) and untreated plants (UNT). The synthetic peptide used was obtained as previously described [13]. The total amount of spray across all the treatments combined was 13 mL per plant, achieving comprehensive coverage of each plant aerial part at every exogenous application. Among a total of 75 tomato plants, 30 were used for each of the G and CTRL experimental groups, and 15 were used for the UNT group. Each replicate was generated by pooling 12 leaves from three individual plants. For the downstream analyses, leaf samples were collected in June 2023, one day following the last spray application.

### Leaf sample processing and microbial DNA extraction

Leaf microbial communities were collected following Gupta et al. [20], with minor modifications. Using ethanol-sterilized scissors, five leaflets from each of 12 leaves per replicate were placed in sterile Ziplock bags and kept on ice until further laboratory processing. To isolate epiphytic microbes, 240 mL of 0.1 M sterilized potassium phosphate buffer (PPB, pH 8) was added, followed by gentle manual shaking, 5 min sonication (50 kHz, Falc Instruments, Treviglio, Italy), and 30 s vortexing. This process was repeated twice. The resulting washing mixture was centrifuged at 11,000 *× g* for 20 min at 4°C, and collected microbial pellets were resuspended in PPB, transferred into 2 mL tubes, and centrifuged at 14,000 *× g* for 2 min at 4°C. Pellets were stored at –20°C until DNA extraction, which was performed using DNeasy PowerSoil Pro Kit (Qiagen, Hilden, Germany) according to the manufacturer’s instructions and quantified with the Qubit HS Assay (Thermo Fisher Scientific, Waltham, Massachusetts, United States).

### Quantitative Real-Time PCR (qPCR) analysis of bacterial abundance

Total bacterial abundance and selected taxa were quantified using SYBR Green-based qPCR with universal primers 515f–806r [103], and taxon-specific primers for Alphaproteobacteria (ALF28f/ALF986r), Gammaproteobacteria (Gamma395f/Gamma871r), Actinobacteria (243f-513r), and *Sphingomonas* (Sph-spt694f-Sph-spt983r) [104–107], amplifying a gene for serine palmitoyltransferase (*spt*). Reaction mixtures contained 1 μL of extracted DNA, 5 μL KAPA SYBR® FAST qPCR Master Mix 2X (KAPA Biosystems, USA), 1 μL of each 10 μM primer, and 3 μL ultrapure water. Amplification was performed on a Q-Tower 3 (Analytik Jena, Germany), with an initial denaturation step at 95 °C for 10 min, followed by 40 cycles of denaturation at 95 °C for 30 s, primer-specific annealing for 30 s at 54 °C (total bacteria, Alpha-, and Gammaproteobacteria), 63° (Actinobacteria), or 55 °C (*Sphingomonas*), extension at 72 °C for 30 s, concluding with a final melting curve analysis. Differences in the abundance of bacteria, among the three different conditions (UNT, CTRL, G) were evaluated using Dunn’s test on log_10_ transformed bacterial gene copy number data.

### Shotgun metagenomic sequencing and analysis

Shotgun metagenomic sequencing was performed by the sequencing provider Novogene Co., Ltd. (Beijing, China). Libraries were prepared using the Nextera XT Index Kit v2 (Illumina, San Diego, California, United States), and sequencing was performed on an Illumina NovaSeq platform (Novogene Europe), leading to 2 × 150 bp reads. Of the 25 samples submitted for sequencing, 22 were successfully processed, resulting in 10 samples for the G treatment, 8 for the CTRL group, and 4 for the UNT group. Quality control of raw reads involved adaptor trimming and quality filtering (Phred < 20) using Trimmomatic v0.39 [108] and VSEARCH v2.15.2 [109]. Host DNA contamination was removed by aligning reads to the *Solanum lycopersicum* reference genome (GCF_000188115.5) using Bowtie 2 [110]. Unmapped reads were assembled with MEGAHIT v1.2.9 [111], retaining contigs >1 kb.

Taxonomic classification and species abundance estimation were conducted using Kraken2 v2.0.9 and Bracken v2.6.0 [112, 113]. Open reading frames (ORFs) were predicted with Prodigal v2.6.3 [114], and a non-redundant, protein-coding gene catalog was built using CD-HIT-EST v4.8.1 [115] (nucleotide identity cutoff 95%). Gene function and taxonomy were assigned using DIAMOND and eggNOG-mapper [116, 117] against the eggNOG v5.0 database [118]. Gene abundances were estimated by mapping back quality-filtered reads to the non-redundant, protein-coding gene catalog using BWA v0.7.17 and SamTools v1.7 [119, 120].

### Binning of bacterial metagenome-assembled genomes (MAGs) and PGPB trait prediction

MAGs were reconstructed using Maxbin2 v2.2.7, MetaBAT2 v2.12.1 and CONCOCT v1.1.0 assembly tools [121–123]. Quality assessment was performed using CheckM v1.0.13 [124], retaining medium-quality bins (completeness >50%, contamination <10%), then dereplicated via both DASTool v1.1.1 [125] and DRep [126]. Taxonomic classification and functional annotation were conducted with GTDB-Tk [127] and DRAM [128]. Abundance profiles for each MAG were determined using CoverM v0.4.0 with –rpkm mode [129] while phylogenetic relationships among MAGs were analyzed using PhyloPhlAn database [130] to construct a phylogenetic tree. Protein FASTA sequences of dereplicated, high-quality MAGs (completeness >90% and contamination <5%) from selected bacterial species were further used for predicting plant growth-promoting traits with the PLaBAse PGPT-Pred tool [43] in strict mode (BLASTP + HMMR).

### Characterization of bacterial diversity and composition

Alpha diversity analysis was performed by normalizing the abundance data to the lowest number of reads to adjust for unequal sequencing depth. The abundance data were normalized using MetagenomeSeq’s cumulative sum scaling (CSS) [131] and subsequently used for beta diversity analysis. The microbial datasets were analyzed with the R packages Phyloseq, MicrobiomeAnalyst, and vegan, all implemented in RStudio [132–135]. Significant differences in alpha diversity metrics (observed species, Shannon index, and Pielou’s evenness) were assessed via the Kruskal‒Wallis test as a nonparametric approach. Differences in community composition between groups were investigated using a normalized Bray‒Curtis dissimilarity matrix and further evaluated by a Permutational multivariate analysis of variance (PERMANOVA) using adonis2 function implemented in the vegan package [134, 135]. Differential abundance of bacterial taxa or genes in metagenomes were explored using the EdgeR package v4.2.0 [136, 137]. Results were considered significant at P_adjusted_ value < 0.05, filtered for log_2_-fold change (log_2_FC) > 2 for enrichment and log_2_FC <-1 for depletion, and with a relative abundance of bacterial species > 0.05%.

### Co-occurrence bacterial network and functional analysis of pathways

Bacterial co-occurrence networks were built using the ggClusterNet package [138], based on Spearman correlations (*r* > 0.7 for positive correlations and *r* < –0.7 for negative correlation), with *P* < 0.05. The resulting correlation matrices were then further analyzed and visualized with Gephi software [139].

Functional enrichment analysis of KEGG Orthology (KO) terms was conducted using Microbiome Profiler v1.10.0 package [140], considering significantly enriched (log_2_FC > 2, FDR < 0.05) or depleted (log_2_FC < –1, FDR < 0.05) terms between CTRL condition and G-treated samples. KO pathway analysis was performed using KEGGREST package v1.44.1 in RStudio [141].

## Ethics approval and consent to participate

Not applicable.

## Consent for publication

Not applicable.

## Availability of data and materials

Phyllosphere metagenomes are available in the European Nucleotide Archive (ENA) (http://www.ebi.ac.uk/ena) under the project number PRJEB86145. Shotgun metagenome reads were deposited under accession numbers ERS23746230-ERS23746251.

All the data generated or analyzed during this study are included in this published article and its supplementary information files.

## Authors’ contribution

Study conception and design: VC and RR. Experimental work: VC and MCC. Methods or reagents provided by: FDF. Analytic and computational tools provided by: WAW and GB. Data analysis: VC and WAW. Interpretation of results: VC, WAW, and GB. Manuscript writing: VC and RR, with contributions from WAW, GB, and FDF. All authors have read and approved the final manuscript.

## List of abbreviations

ProSys: Prosystemin
UNT: untreated leaves
CTRL: PBS 0.1X treated leaves
G: Prosystemin-derived peptide
KO: KEGG Orthology

## Acknowledgments

We sincerely thank Dr. Anna Aprile (University of Naples “Federico II”) for helping in samples collection and processing; Anja Lamprecht (Graz University of Technology) for quantitative real-time PCR experiment; Materias S.r.l. for encouragement and support.

**Description of supplemental materials**

**Additional file 1:**

**Table S1: Number of sequencing reads per sample.**

**Table S2: Differential abundance analysis between peptide (G) and PBS0.1X-treated (CTRL) tomato phyllosphere samples.**

**Table S3: Shared core and unique core bacterial species.**

**Table S4: KEGG_KO enrichment analysis of bacterial genes associated with G peptide treatment.**

**Table S5: Topological properties of bacterial networks.**

**Table S6: differentially enriched bacterial genes associated to two-component system (TCS) pathway in G-treated samples compared to CTRL condition.**

**Table S7: CheckM results with medium-quality assembled genomes (MAGs) and relative taxonomy classification.**

**Table S8: Number of mapped Reads Per Kilobase per Million reads (RKPM) for each Metagenome-Assembled Genome (MAG) across the different treated samples.**

**Table S9: PLaBAse annotation.**

**Additional file 2:**

**Figure S1. Two-component system overview of the tomato phyllosphere microbiome.**

**Figure S2. Phylogenetic tree and abundance profiles of metagenome-assembled genomes (MAGs) across different treatment of tomato leaves.**

