## Additional file 2 for "Prosystemin–derived signals: bridging leaf microbiome dynamics and defense activation"

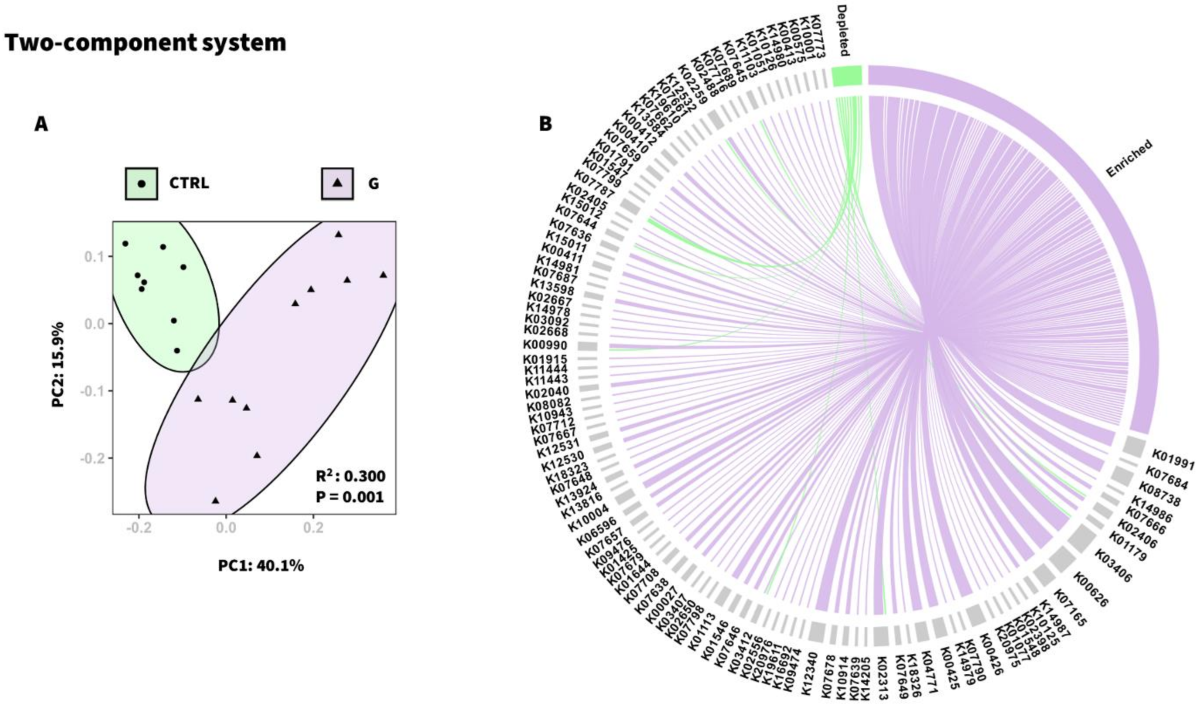


**Figure S1. Two-component system overview of the tomato phyllosphere microbiome.** PCoA based on Bray–Curtis metrics (**A**) showing differences between CTRL and G-treated samples on the basis of the TCS-associated genes. Chord diagram (**B**) showing the connections between the source represented by genes classified as enriched or depleted in the peptide-treated samples ($log_2FC > 2$ or $< -1$; $FDR < 0.05$; EdgeR) and the targets represented by associated KEGG terms related to the TCS pathway (B). Enriched or depleted KEGG terms are depicted in violet or green, respectively. The thickness of the chords represents the $log_2FC$ values, which range from -1 to 7.92.

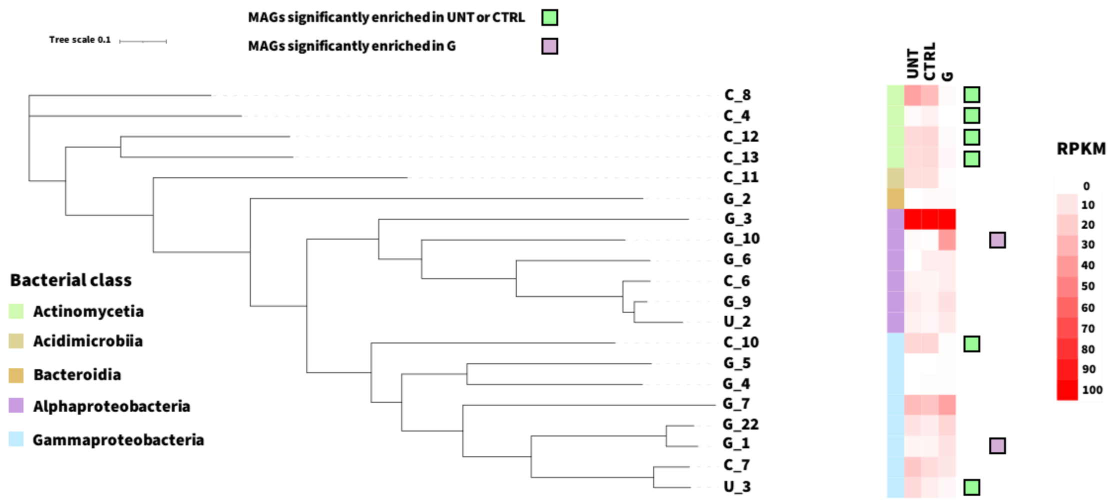


**Figure S2. Phylogenetic tree and abundance profiles of metagenome-assembled genomes (MAGs) across different treatment of tomato leaves.** Phylogenetic tree of dereplicated MAGs (completeness >50% and contamination <10%), coloured according to bacterial class taxonomy. Abundance of Reads Per Kilobase per Million mapped reads (RPKM) values for each MAG across untreated (UNT), control (CTRL), or peptide-treated (G) samples, are reported in the right panel. Significantly enriched MAGs are represented by green squares for control conditions (UNT/CTRL) or violet squares for G peptide treatment.
